# β-nicotinamide mononucleotide production in *Vibrio natriegens*: a preliminary study

**DOI:** 10.1101/2025.03.17.643733

**Authors:** Ly Huong Tran, Vu Thao Phuong Tran, Huy Tuan Dat Pham, Giang Tra Nguyen, Tuoi Thi Nghiem, Anh Duc Pham, Nga Quynh Pham, Thuy Thi Le, Ngoc Lan Nguyen, Tran Nhat Minh Dang, Thinh Huy Tran, Van Khanh Tran, Hoa Quang Le

## Abstract

**Background:** Nicotinamide mononucleotide (NMN), is a promising nutraceutical attracting much attention for its pharmacological and anti-aging efficacies. However, NMN-containing commercial products are very high-priced due to the lack of efficient and facile methods for industrial-scale production. To date, various metabolic engineering strategies have been successfully applied to produce NMN in *Escherichia coli*. Recently, *Vibrio natriegens* has become a promising host in the bioindustry thanks to its rapid growth and capabilities of broad substrate utilization. This study aims to evaluate the NMN biosynthesis capability of *V. natriegens*.

**Methods:** Firstly, a mutant *V. natriegens* strain *(Δdns::araC-T7RNAP-Kan*^*R*^ *ΔpncC::FtnadE-Sm*^*R*^ *ΔnadR)* was generated via multiplex genome editing by natural transformation (MuGENT). *Nampt* genes encoding nicotinamide phosphoribosyltransferase from *Chitinophaga pinensis, Sphingopyxis sp*. C-1, *Haemophilus ducreyi, and Vibrio* phage KVP40 were codon-optimized and cloned into pACYCDuet™-1 under the control of T7 promoter. The recombinant plasmids were electroporated into the mutant strain. The expression of recombinant NAMPTs in *V. natriegens* was evaluated by SDS-PAGE analysis and the intracellular NMN concentrations were quantified by HPLC.

**Results:** After two rounds of MuGENT, *V. natriegens* V54-33 strain *(Δdns::araC-T7RNAP-Kan*^*R*^ *ΔpncC::FtnadE-Sm*^*R*^ *ΔnadR)* was successfully generated. SDS-PAGE analysis demonstrated that all NAMPTs were strongly expressed in the V54-33 strain. HPLC analysis revealed that the highest intracellular NMN concentration was obtained with NAMPT from *Chitinophaga pinensis* (44.5 μM), followed by NAMPT from *Vibrio* phage KVP40 (23.3 μM).

**Conclusion:** This study demonstrated the feasibility of NMN biosynthesis in *V. natriegens*.

## INTRODUCTION

β-nicotinamide mononucleotide (NMN) is a precursor of NAD^+^, an essential coenzyme in various metabolic processes. NAD^+^ is involved in intracellular signaling pathways, regulating mitochondrial functions and other biochemical processes.^1^ In living organisms, intracellular NAD^+^ levels are directly influenced by its precursor, NMN. Recent studies reported various potential medical applications of NMN, particularly in anti-aging, cognitive-enhancing, and the treatment of severe chronic diseases such as Alzheimer’s and diabetes.^2,3^

There are three major ways for NMN production: chemical synthesis, enzymatic methods and fermentation methods using genetically modified microbial strains.^4^ The chemical synthesis pathway offers the advantage of low production costs; however, the resulting isomeric compounds may exhibit toxicity or lack of biological activity. The enzymatic methods enhance selectivity and reduce toxicity, but the high costs of enzymes and phosphate sources such as ATP remain a major obstacle. Recent studies indicated that the fermentation approach using metabolically engineered microbial strains represents a promising solution to overcome the limitations of the latter methods. The most effective strategy for NMN biosynthesis in microbial cells primarily relies on: nicotinamide (NAM) supplement as a substrate, activation of pentose phosphate pathway to generate the precursor PRPP from carbon sources, overexpression of nicotinamide phosphoribosyltransferase (NAMPT) to catalyze the reaction between NAM and PRPP to produce NMN, and the expression of NMN transporter.^5,6^ Among these factors, NAMPT is the key enzyme that dictates NMN production efficiency.^5–7^ In a pioneer study, Marinescu *et al*. evaluated NAMPT from various sources and found that NAMPT from *Haemophilus ducreyi*, when expressed in *E. coli* BL21(DE3)pLysS, gave the highest intracellular NMN concentration (15.42 mg/L).^7^ Subsequently, Black *et al*. has combined the overexpression of both NAMPT and NMN synthase from *Francisella tularensis* (FtNadE) with the deletion of *pncC* and *nadR*, two genes involved in the catabolism of NMN, in *E. coli*, and achieved an intracellular NMN concentration of 1.5 mM.^8^ In another study, NAMPTs from ten organisms have been tested for the activity when expressed in *E. coli* and the results revealed that NAMPTs from *Chitinophaga pinensis* and *Sphingopyxis* sp. C-1 displayed significantly higher activities compared to NAMPTs derived from other sources.^5^ Similarly, Huang *et al*. performed a screening of eight NAMPTs from various sources and found that NAMPT from *Vibrio* phage KVP40 could biosynthesize 81.3 mg/L NMN intracellularly.^6^

Recently, *Vibrio natriegens* has emerged as a host for industrial biotechnology due to its extremely rapid growth rate with a reported doubling time of less than 10 min, non-pathogenic nature, genetic tractability, and ability to utilize a variety of carbon sources.^9^ In numerous studies, *V. natriegens* was used as an alternative expression system for recombinant protein production,^10^ and high-value metabolites such as pyruvate,^11^ 2,3-butanediol.^12^ However, the application of *V. natriegens* in NMN production remains unexplored. Therefore, the present study aimed to evaluate the feasibility of NMN biosynthesis in *V. natriegens*.

## MATERIALS AND METHODS

### Materials

*Vibrio natriegens* strain TND1964, kindly offered by Prof. Ankur Dalia from Indiana University, Bloomington, USA and *Escherichia coli* DH5α (Thermo Scientific™, cat. number EC0112) were used as hosts for expression and cloning, respectively.

Oligonucleotides were synthesized by Macrogen (Korea). Expression vector used in this study was pACYCDuet™-1 (Novagen, cat. number 71147-3).

All other reagents were obtained from Thermo Scientific, New England Biolabs, Merck, Qiagen, and Bio Basic, unless otherwise specified.

### Generation of the recombinant *Vibrio natriegens* V54-33 strain

*Vibrio natriegens* V54-33 strain *(Δdns::araC-T7RNAP-Kan*^*R*^ *ΔpncC::FtnadE-Sm*^*R*^ *ΔnadR)* was generated using multiplex genome editing by natural transformation (MuGENT) method.^13^ Details were presented in the Supplement I.^14^

### Construction of the *pACYC-nampt* plasmids

The *Nampt* genes encoding nicotinamide phosphoribosyltransferase (NAMPT) from *C. pinensis, Sphingopyxis* sp. C-1, *H. ducreyi*, and *Vibrio* phage KVP40, were codon-optimized for expression in *V. natriegens* and synthesized by Genscript. These genes were cloned into the pACYCDuet™-1 (Novagen, cat. number 71147) under the control of T7 promoter-1 (Supplement II).^14^

### Electroporation into *Vibrio natriegens* V54-33 strain

The preparation of *V. natriegens* electrocompetent cells and the electroporation of DNA plasmids were carried out following the procedures described by Weinstock *et al*.^15^ Transformants were selected on LBv2 agar plates supplemented with Chloramphenicol (25 μg/mL) (Bio Basic, cat. number CB0118). LBv2 is LB-Miller (Bio Basic, cat. number SD7003) supplemented with v2 salts (204 mM NaCl (Bio Basic, cat. number DB0483), 4.2 mM KCl (Bio Basic, cat. number PB0440, and 23.14 mM MgCl_2_ (Bio Basic, cat. number MB0328)).

### Shake Flask Fermentation

A single colony was inoculated into 2 mL of LBv2 medium supplemented with Chloramphenicol (25 μg/mL) (Bio Basic, cat. number CB0118) and cultured at 30 °C, 200 rpm overnight. Subsequently, 1% (v/v) of the overnight culture was reinoculated into 10 mL of fresh LBv2 at 30 °C and 200 rpm. Induction was carried out by adding 2 g/L arabinose (Bio Basic, cat. number AB0071L) when OD_600_ reached 0.6−0.8. The cultures were cultivated until OD_600_ reached 5-6 and the cultured cells, corresponding to 5 mL cultures, were collected for SDS-PAGE analysis. The remaining cultures were centrifuged at 3000 *g* for 5 min at 4 ^o^C. After discarding the supernatant, the cells were washed in 20 mL modified M9 medium^16^ and resuspended in 5 mL modified M9 medium supplemented with Chloramphenicol (25 μg/mL) (Bio Basic, cat. number CB0118), 1 mM nicotinamide (Sigma, cat. number N0636) and 1 mM nicotinic acid (Sigma, cat. number N0765). The cells were cultured at 30 ^**o**^C and 200 rpm for 5 h and cultured cells, corresponding to 1 OD_600_, were collected, washed with 1 mL of water and stored at -80 ^o^C until NMN quantification.

### NMN quantification by HPLC

NMN quantification was carried out according to the method described by Vu *et al*.^17^ Briefly, cells were resuspended in 500 μL of water and lysed by sonication on ice by using VC 130 System (Sonics & Materials, Inc, cat. number VC130). The amplitude was set at 80% and on/off pulse of 10 s duration each was given for 1 min. Subsequently, 100 μL of trichloroacetic acid 25% (Bio Basic, cat. number TB0968) was added to inactivate the enzymes in the lysate. The obtained mixture was clarified by centrifugation at 15000 *g* for 15 min at 4 ^o^C and the supernatant was used for HPLC analysis. Supernatants were run on a Shimadzu LC-20A high-performance liquid chromatography (HPLC) system with a Reliant^TM^ C18 chromatography column (150 mm x 4.6 mm, 5 µm) (Waters, cat. number 186007282). Mobile phase used for separation was 10 mM phosphate buffer pH 3: methanol (90:10) (Supelco, cat. number 106007), at the flow rate of 1.0 mL/min. The injection volume was 20 μL, and individual peak areas were determined using a photo diode array at 261 nm.

## RESULTS AND DISCUSSION

In our present study, a metabolic engineering approach (Figure 1) combining the insertion of key enzymes in the NMN biosynthetic pathway (NMN synthase from *Francisella tularensis* (FtNadE) and the most promising NAMPTs from previous studies) and the inactivation of endogenous NMN-degrading enzymes (NMN amidohydrolase, encoded by *pncC*, and NMN adenylyltransferase, encoded by *nadR*) was used to evaluate the NMN production capability in *V. natriegens*. To this end, a mutant strain, called V54-33, has been successfully generated by two rounds of MuGENT introducing araC-T7 RNA polymerase construct and *FtNadE* under the control of T7 promoter, respectively, while deleting *pncC* and *nadR* genes (Supplement I).^14^ This allowed a tight induction of NAMPTs and FtNadE by arabinose, and thereby provided a controlled regulation for NMN biosynthesis. Of note, our previous attempts using T7 IPTG-induced system were unsuccessful because of the growth inhibition in transformants (possibly due to the leaky expression of T7lac promoter) (data not shown).

**Figure 1.**
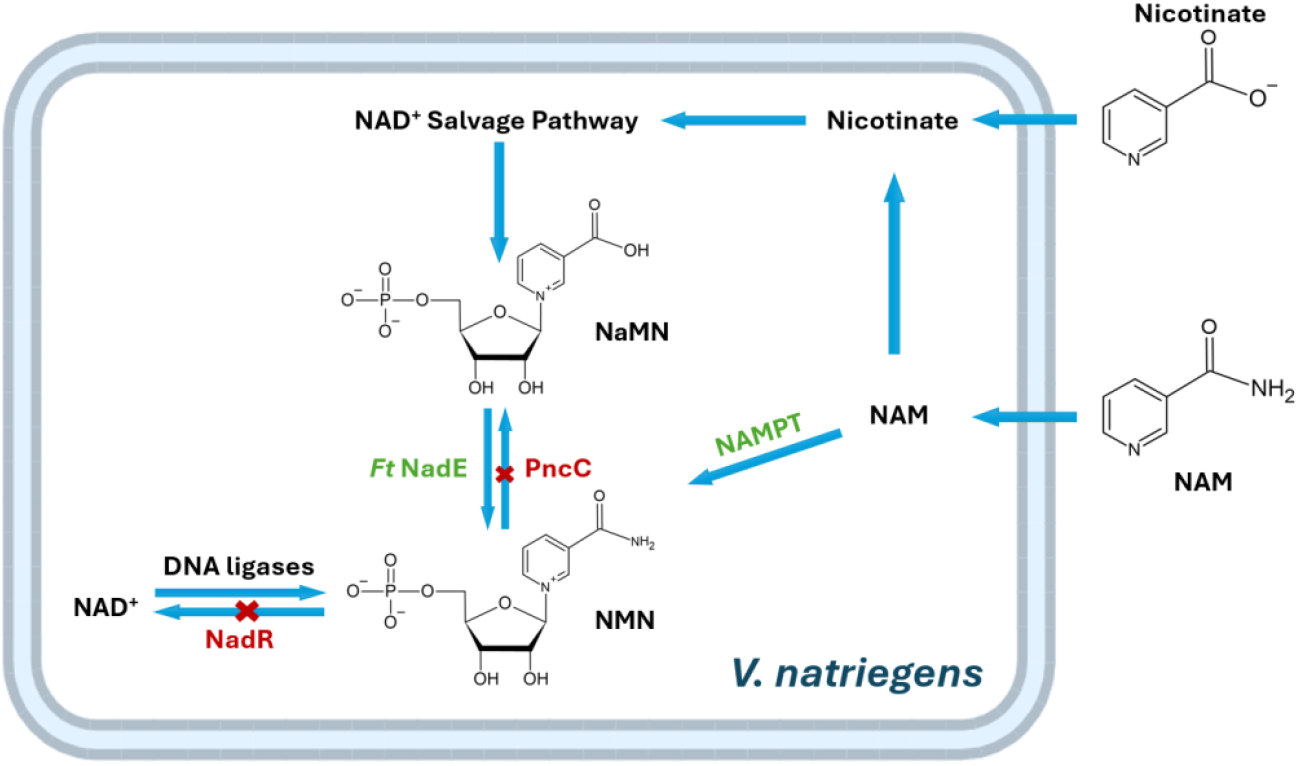
Biosynthesis routes of β-nicotinamide mononucleotide used in this study (modified from 8). NMN: nicotinamide mononucleotide, NaMN: nicotinic acid mononucleotide, NAD+: nicotinamide adenine dinucleotide, NAM: nicotinamide, NAMPT: nicotinamide phosphoribosyltransferase, PncC: NMN amidohydrolase, FtNadE: NMN synthetase from *Francisella tularensis*, NadR: NMN adenylyltransferase.

Subsequently, four plasmids overexpressing NAMPTs from *C. pinensis* (CP), *Sphingopyxis* sp. C-1 (SSC), *H. ducreyi* (HD) and *Vibrio* phage KVP40 (VP) under the control of T7 promoter, were successfully generated (Supplement II).^14^ These four enzymes have been reported as the most efficient NAMPTs in separated studies.^5–7^ In the next step, the strain V54-33 has been transformed with these plasmids harboring NAMPTs. SDS-PAGE analysis (Figure 2) clearly showed that all recombinant strains carrying NAMPT plasmids displayed an intense band at the expected size of 56 kDa after induction with arabinose, while the original V54-33 strain did not possess this band. These results demonstrated that NAMPTs from CP, HD, SSC and VP were successfully expressed in *V. natriegens*.

**Figure 2.**
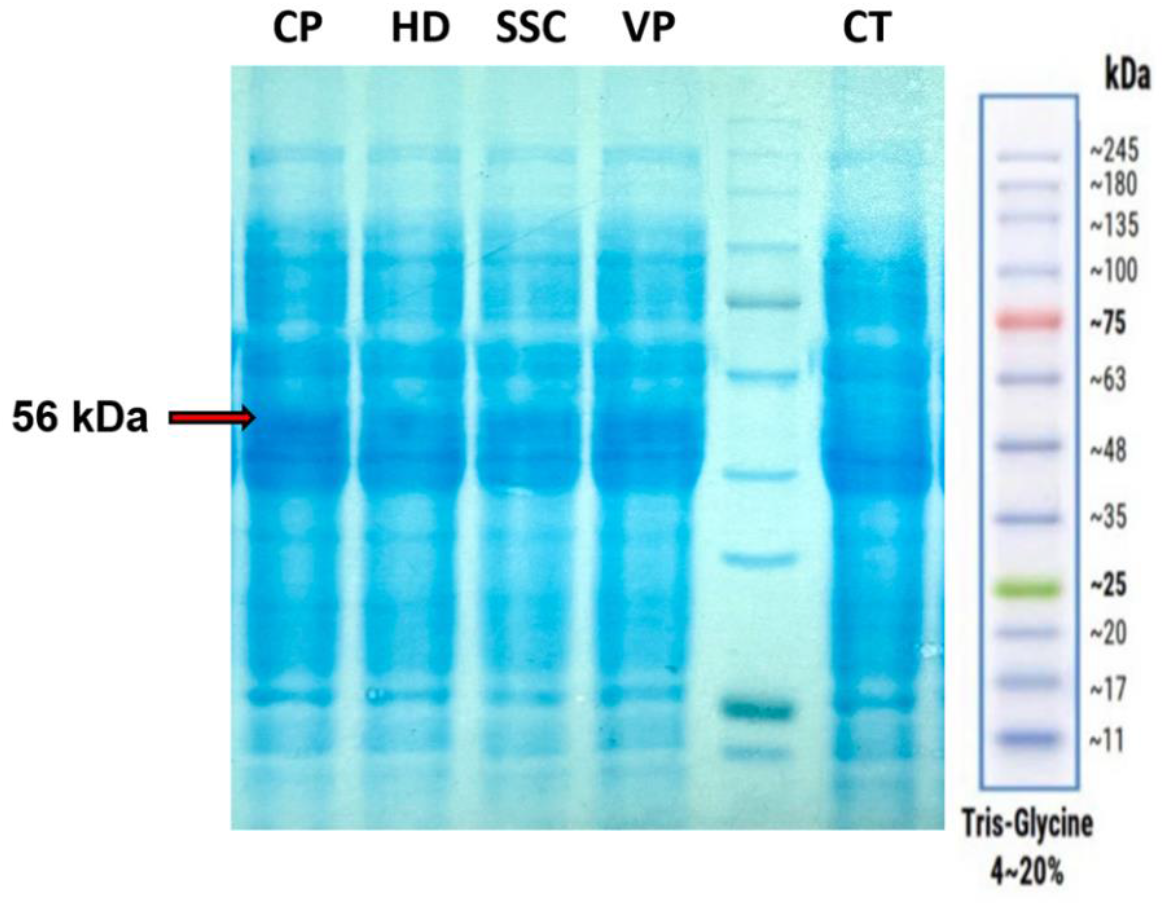
SDS-PAGE analysis of the expression of recombinant NAMPTs in V54-33 strain. L: GangNam-STAIN™ Prestained Protein Ladder. CP: lysate of V54-33 strain overexpressing NAMPT from *Chitinophaga pinensis*. HD: lysate of V54-33 strain overexpressing NAMPT from *Haemophilus ducreyi*. SSC: lysate of V54-33 strain overexpressing NAMPT from *Sphingopyxis* sp. C-1. VP: lysate of V54-33 strain overexpressing NAMPT from *Vibrio* phage KVP40. CT: lysate of V54-33 strain.

These recombinant strains were then transferred into modified M9 medium to assess NMN biosynthesis. HPLC quantification revealed that three out of four transformants (CP, HD, and VP) were capable of intracellular NMN accumulation, while SSC transformants and the V54-33 strain exhibited NMN levels below the limit of detection (LOD) of the method (Figure 3, Supplement III).^14^ Among them, CP transformants exhibited the highest intracellular NMN titers (44.5 μM), whereas HD transformants showed significantly lower NMN accumulation (12.1 μM, p = 0.0111) (Figure 3, Supplement III).^14^ The NMN concentration of VP transformants was intermediate (23.3 μM) (Supplement III),^14^ which aligns with prior research by Huang *et al*., showing that NAMPT from VP was more efficient in NMN accumulation than NAMPT from HD.^6^ When considering the absolute NMN titers reported in *E. coli*, NMN accumulation in CP transformants was comparable to that reported by Marinescu *et al*. (15.42 mg/L),^7^ but significantly lower than those reported by Black *et al*. (1.5 mM)^8^ and Huang *et al*. (81.3 mg/L).^6^ These disparities could be explained by the difference in expression systems: (i) *FtNadE* was expressed on a multiple-copy plasmid in *E. coli* in the study by Black *et al*.^*8*^ while this gene was incorporated into the chromosome of *V. natriegens* as a single-copy in our study; (ii) *ushA* encoding 5′-phosphatase capable of dephosphorylation of NMN to nicotinamide riboside and *purR* regulating the synthesis of the precursor phosphoribosyl pyrophosphate (PRPP) were knocked out in the study of Huang *et al*.^*6*^ whereas these genes have not been inactivated in the present study.

**Figure 3.**
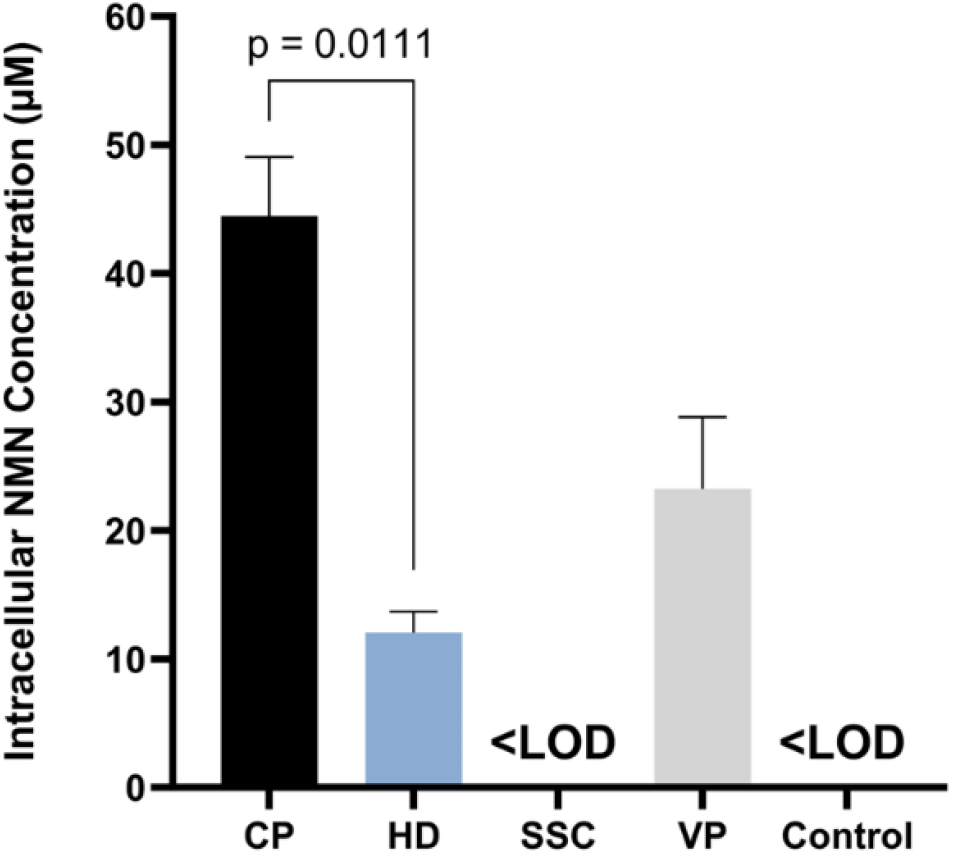
Quantification of intracellular NMN concentration of V54-33 strains harboring pACYC-Nampt plasmids. CP: V54-33 strain overexpressing NAMPT from *Chitinophaga pinensis*. HD: V54-33 strain overexpressing NAMPT from *Haemophilus ducreyi*. SSC: V54-33 strain overexpressing NAMPT from *Sphingopyxis* sp. C-1. VP: V54-33 strain overexpressing NAMPT from *Vibrio* phage KVP40. Control: V54-33 strain. <LOD: concentration lower than limit of detection of HPLC method. Data are expressed as mean ± S.D. (error bars). Statistical analysis of differences between groups was performed by unpaired t-test (p < 0.05).

In summary, this study demonstrated the feasibility of NMN biosynthesis in *V. natriegens*. Further research focusing on the activation of pentose phosphate pathway and the transporters of NMN and NAM is underway to improve NMN production in this promising host.

## Supporting information

Supplement III

## ETHICS AND CONSENT

Ethical approval and consent were not required.

## DATA AVAILABILITY

### Underlying data

Dataset from NCBI Sequence Read Archive: Sequencing results for bioproject: “β-nicotinamide mononucleotide production in Vibrio natriegens: a preliminary study”

Accession number PRJNA1231393; https://www.ncbi.nlm.nih.gov/sra/?term=PRJNA1231393

Data are available under the terms of the Creative Commons Zero “No rights reserved” data waiver (CC0 1.0 Public domain dedication).

### Extended data

Figshare: Supplement for “β-nicotinamide mononucleotide production in Vibrio natriegens: a preliminary study”.^14^

https://figshare.com/articles/dataset/Supplement_for_-nicotinamide_mononucleotide_production_in_Vibrio_natriegens_a_preliminary_study_/28547603/1

This project contains the following extended data:

- Supplement I: Generation of *Vibrio natriegens* strain V54-33 *(Δdns::araC-T7RNAP-Kan*^*R*^ *ΔpncC::FtnadE-Sm*^*R*^ *ΔnadR)*.pdf
- Supplement II: Generation of pACYCDuet-Nampt plasmids.pdf
- Supplement III: Quantification of intracellular NMN concentration by HPLC.pdf

Data are available under the terms of the Creative Commons Zero “No rights reserved” data waiver (CC0 1.0 Public domain dedication).

