## Supplement I for "β-nicotinamide mononucleotide production in *Vibrio natriegens*: a preliminary study"

### Generation of *Vibrio natriegens* strain V54-33 (*Δdns::araC-T7RNAP-Kan$^R$ ΔpncC::FtnadE-Sm$^R$ ΔnadR)*

**Supplement Table I.1. Primers used for generation of *Vibrio natriegens* strain V54-33.**

| Primer name | Primer Sequence (5'→3') | Description |
|---|---|---|
| **Primers for mutant constructs** | | |
| Vna-deldns-F1 | TGTACTAGCAAGCGTGGTATTCAGCA | *Δdns* F1 |
| Vna-deldns-R1 | GGCACTGGATAGTGCAAGAATGA | *Δdns* R1 |
| Pbad-araC-gb-F | ttcttgcactatccagtgcc TGTCAAATGGACGAAG | *araC* F |
| Pbad-araC-gb-R | AGAAACAGTAGAGAGTTGCGA | *araC R* |
| T7pol-gb-F | tcgcaactctctactgtttct ACTTTAATAAGGAGATATACCATGA | T7RNAP F |
| T7pol-gb-R | AGCATCCTCCTTAATTCAAGATCC TTACGCGAACGCGAAGTC | T7RNAP R |
| KanR-gb-F | GGATCTTGAATTAAGGAGGATGCT AGCTAATACGACTCACTATAGG | Kan$^R$ F |
| KanR-gb-R | tgagtctgttcgacttaagcatta ATTCTTAGAAAAACTCATCGAGCA | Kan$^R$ R |
| Vna-deldns-F2 | taatgcttaagtcgaacagactca CCAATCGCGACAATCG | *Δdns* F2 |
| Vna-deldns-R2 | GCTTCAAGCATCATGGCAAAGCTAGA | *Δdns* R2 |
| Vna-delpncC-F1 | TGCGTTTTAAGCAGGGTATCCA | *ΔpncC* F1 |
| Vna-delpncC-R1 | tcccctatagtgagtcgtattaatt CTGCTCGAGTAACACACCT | *ΔpncC* R1 |
| T7pro-FtNadE-gb-F | aattaatacgactcactataggggа GTACATTAAAGAGGAGAAATATACAT | $P_{T7}$-*FtnadE* F |
| FtNadE-gb-R | gtcgctctgttcgacttaagcatta GAAGTTCGGAGTTAGAG | $P_{T7}$-*FtnadE* R |
| SmR-gb-F | taatgcttaagtcgaacag AGCGACCGAGTGAGCTAG | Sm$^R$ F |
| SmR-gb-R | atatcgttatgaaatcccaattatt TGCCGACTACCTTGGT | Sm$^R$ R |
| Vna-delpncC-F2 | aataattgggatttcataacgatat GGCAGCACTGACTCCAA | *ΔpncC* F2 |
| Vna-delpncC-R2 | ACCAGCAGTACCCATGACT | *ΔpncC* R2 |
| Vna-delnadR-F1 | CACCGCGACAGGTGGAAGA | *ΔnadR* F1 |
| Vna-delnadR-R1 | TGTGGCCTAAGCTAGGTGGA | *ΔnadR* R1 |

| | | |
|---|---|---|
| Vna-delnadR-F2 | aatccacctagcttaggccac ACTGAGCGCTGGACCGT | *ΔnadR* F2 |
| Vna-delnadR-R2 | ACGTCTATCATGCGGCTACCGA | *ΔnadR* R2 |
| **Primers for screening transformants** | | |
| T7pol-sc-F | CGTACTCACAGTAAGAAAGCA | Screen for *Δdns::araC-T7RNAP-Kan$^R$* |
| T7pol-sc-R | ACAAGCCATGATGTTCTCGTG | |
| nadR-sc-F | TCATCACGGATGGATTGTTC | Screen for *ΔnadR* |
| nadR-sc-R | GCTACGCTTGCTTGCGA | |
| **Sequencing primers** | | |
| T7pol-seq-F1 | CGCTGGCTGGCATCTCT | T7RNAP sequencing primers |
| T7pol-seq-F2 | ACGGACAACGAAGTAGT | |
| T7pol-seq-R1 | ACAAGCCAGAGTGGTCTTAATGGT | |
| T7pol-seq-R2 | CAGTGCTTTCTTACTGTG | |
| T7pol-seq-R3 | ATCAGGAGTTACCCAATG | |
| FtNadE-seq-F | TCCGGAGCGGCTTAGGA | *NadE* sequencing primer |
| araC-seq-F | ATGAAATACCTGTTCTCTTTATTC | *araC* sequencing primers |
| araC-seq-R | AGAAACAGTAGAGAGTTGCGA | |

**Lower case nucleotides specify overlap regions*

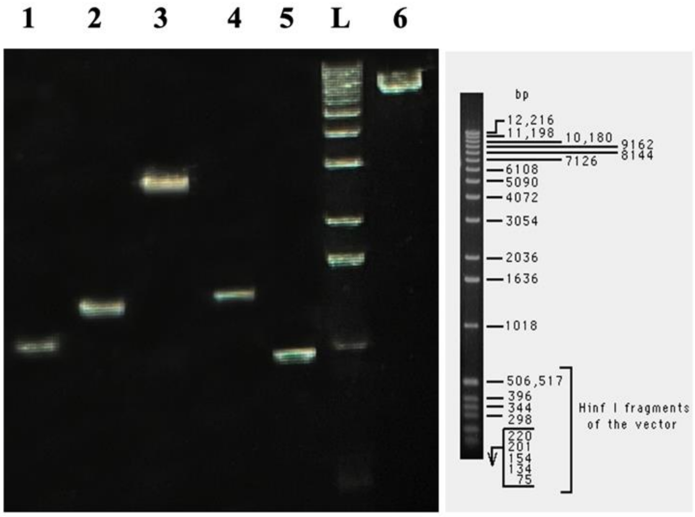


**Supplementary Figure I.1. Agarose gel analysis for verification of PCR products during the preparation of *Δdns::araC-T7RNAP-Kan$^R$* construct by Gibson assembly.**

*(1) flanking left fragment; (2) araC-$P_{BAD}$ fragment; (3) T7RNAP fragment; (4) Kan$^R$ fragment; (5) flanking right fragment; (6) Gibson assembly product amplified by Vna-deldns-F1 and Vna-deldns-R2 primers; L: 1 kb DNA ladder (Invitrogen)*

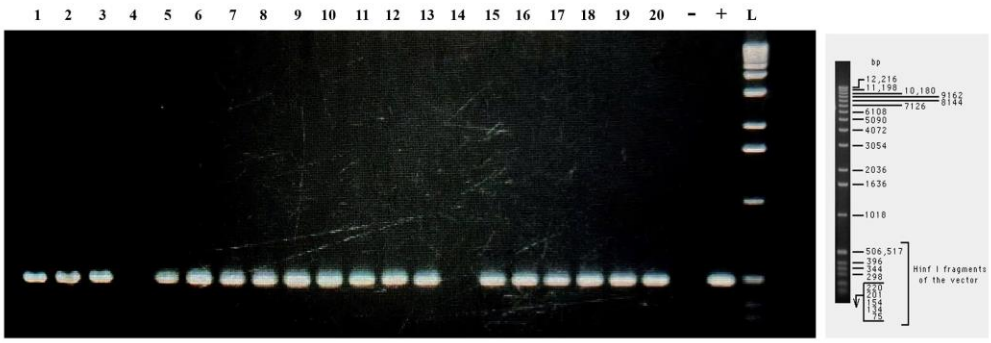


**Supplementary Figure I.2. Screening results for transformants carrying the *Δdns::araC-T7RNAP-Kan$^R$* construct using PCR with T7pol-sc-F and T7pol-sc-R primers.**
*1-20: colony PCR products; (-): no-template control; (+) positive control; L: 1 kb DNA ladder (Invitrogen)*

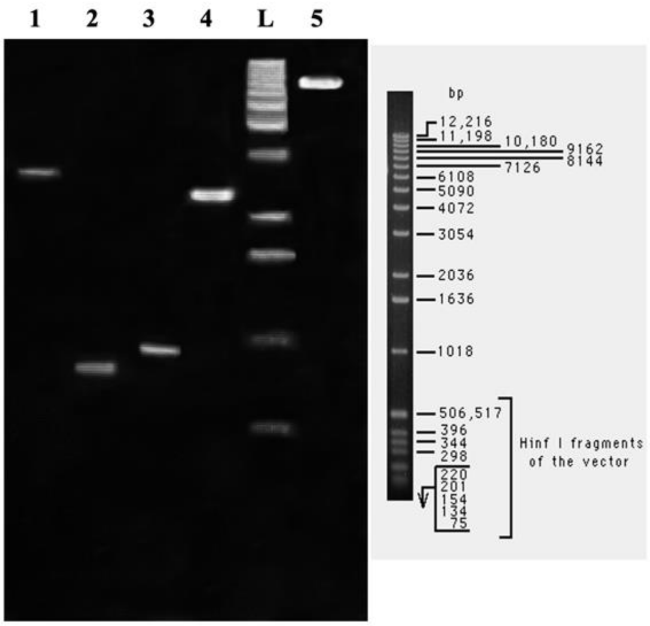


**Supplementary Figure I.3. Agarose gel analysis for verification of PCR products during the preparation of *ΔpncC::FtnadE-Sm$^R$* construct by Gibson assembly.**
*(1) flanking left fragment; (2) FtnadE fragment; (3) Sm$^R$ fragment; (4) flanking right fragment; (5) Gibson assembly product amplified by Vna-delpncC-F1 and Vna-delpncC-R2 primers; L: 1 kb DNA ladder (Invitrogen)*

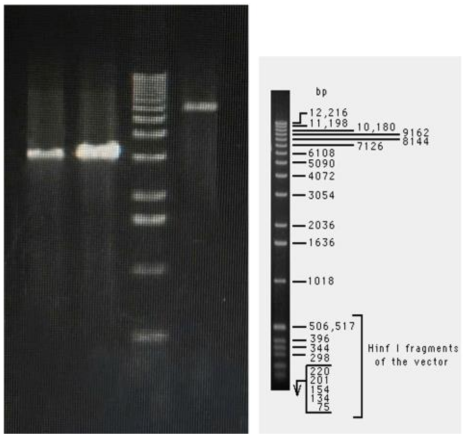


**Supplementary Figure I.4. Agarose gel analysis for verification of PCR products during the preparation of *ΔnadR* construct by Gibson assembly.**

*(1) flanking left fragment; (2) flanking right fragment; (3) Gibson assembly product amplified by Vna-delnadR-F1 and Vna-delnadR-R2 primers; L: 1 kb DNA ladder (Invitrogen)*

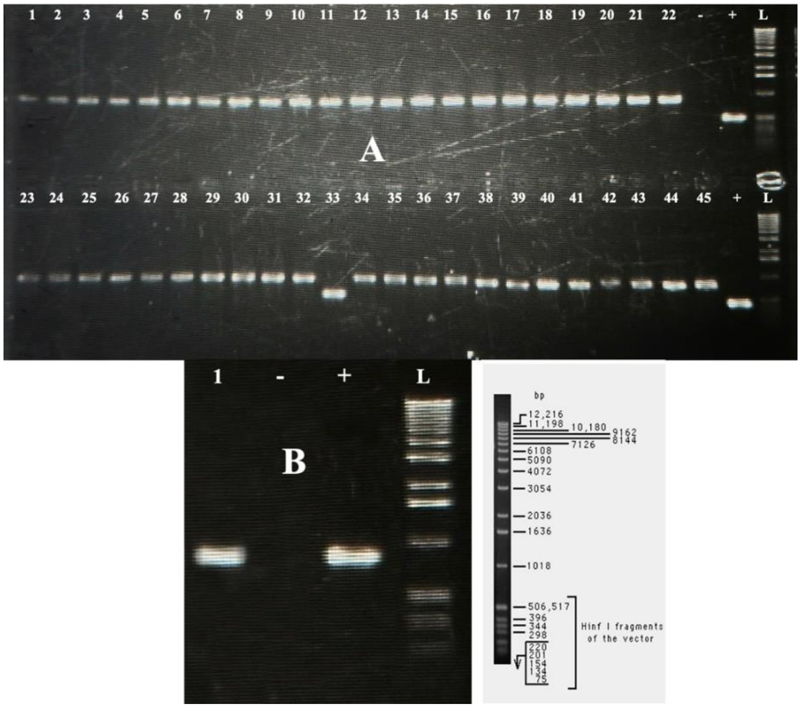


**Supplementary Figure I.5.**

**A. Screening results for mutants with deletion of *nadR* by PCR using nadR-sc-F and nadR-sc-R primers.**

*1-45: colony PCR products; (-): no-template control; (+) positive control; L: 1 kb DNA ladder (Invitrogen)*

**B - Screening results for *ΔpncC::FtnadE-Sm$^R$* construct by PCR using T7pro-FtNadE-gb-F and FtNadE-gb-R primers.**

*1: colony PCR product of strain V54-33; (-): no-template control; (+) positive control; L: 1 kb DNA ladder (Invitrogen)*

In this study, *Vibrio natriegens* strain TND1964, kindly offered by Prof. Ankur Dalia from Indiana University, Bloomington, USA, was genetically modified by introducing linear PCR products via natural transformation following the procedure described by Dalia et al.[1]

Firstly, in order to improve transformation efficiency, the *dns* locus encoding DNA endonuclease from this strain was deleted by introducing the construct araC-$P_{BAD}$-T7RNAP-$Kan^R$. This construct was generated by Gibson assembly from the following fragments:

- the flanking left sequence (1 kb) which was amplified by PCR from *V. natriegens* genomic DNA in a 50 µL reaction containing 0.5 µM of Vna-deldns-F1 and 0.5 µM Vna-deldns-R1, 100 ng of extracted DNA, 0.2 mM dNTPs, and 0.02 U/µL of the Q5 Hot Start High-Fidelity DNA Polymerase in 1X Q5 Reaction Buffer. The thermocycling conditions of the PCR consisted of 98°C for 30 s, followed by 30 cycles of 98°C for 10 s, 68°C for 30 s, and 72°C for 30s.

- *araC-$P_{BAD}$* fragment (1.3 kb) which was produced by PCR amplification in a 50 µL reaction containing 0.5 µM of Pbad-araC-gb-F and 0.5 µM Pbad-araC-gb-R, 100 ng of *E. coli* BL21(DE3) genomic DNA, 0.2 mM dNTPs, and 0.02 U/µL of the Q5 Hot Start High-Fidelity DNA Polymerase in 1X Q5 Reaction Buffer. The thermocycling conditions of the PCR consisted of 98°C for 30 s, followed by 30 cycles of 98°C for 10 s, 64°C for 30 s, and 72°C for 40 s.

- T7RNAP coding sequence (2.7 kb) which was synthesized by Genscript. The fragment was amplified in a 50 µL reaction containing 0.5 µM of T7pol-gb-F and 0.5 µM T7pol-gb-R, 1 ng of template, 0.2 mM dNTPs, and 0.02 U/µL of the Q5 Hot Start High-Fidelity DNA Polymerase in 1X Q5 Reaction Buffer. The thermocycling conditions of the PCR consisted of 98°C for 30 s, followed by 6 cycles of 98°C for 10 s, 59°C for 30 s, 72°C for 1 min 30 s, and 9 cycles of 98°C for 10 s, and 72°C for 2 min.

- $Kan^R$ fragment (1.3 kb) which was produced by PCR amplification in a 50 µL reaction containing 0.5 µM of KanR-gb-F and 0.5 µM KanR-gb-R, 1 ng of modified pET28a vector, 0.2 mM dNTPs, and 0.02 U/µL of the Q5 Hot Start High-Fidelity DNA Polymerase in 1X Q5 Reaction Buffer. The thermocycling conditions of the PCR consisted of 98°C for 30 s, followed by 6 cycles of 98°C for 10 s, 63°C for 30 s, 72°C for 40 s, and 9 cycles of 98°C for 10 s, and 72°C for 1 min 20 s.

- The flanking right sequence (1 kb) which was amplified by PCR from *V. natriegens* genomic DNA in a 50 µL reaction containing 0.5 µM of Vna-deldns-F2 and 0.5 µM Vna-deldns-R2, 100 ng of extracted DNA, 0.2 mM dNTPs, and 0.02 U/µL of the Q5 Hot Start High-Fidelity DNA Polymerase in 1X Q5 Reaction Buffer. The thermocycling conditions of the PCR consisted of 98°C for 30 s, followed by 30 cycles of 98°C for 10 s, 67°C for 30 s, 72°C for 30 s.

Subsequently, all PCR products were verified by agarose gel electrophoresis (Supplementary Figure I.1) and purified by Monarch® PCR & DNA Cleanup Kit (New England BioLabs, cat. number T1030) according to the manufacturer's instructions. These five fragments were then assembled by NEBuilder® HiFi DNA Assembly Master Mix (New England Biolabs, cat. number E2621L) according to the manufacturer's instructions. The resulting product of assembled reaction was amplified in a 50 µL reaction containing 0.5 µM of Vna-deldns-F1 and 0.5 µM Vna-deldns-R2, 1 µL of assembled product, 0.2 mM dNTPs, and 0.02 U/µL of Q5 Hot Start High-

Fidelity DNA Polymerase in 1X Q5 Reaction Buffer. The thermocycling conditions consisted of an initial denaturation at 98°C for 30 s, followed by 15 cycles of 98°C for 10 s, 70°C for 30 s, and 72°C for 4 min. After verification by agarose gel electrophoresis (Supplementary Figure I.1) and purification by Monarch® PCR & DNA Cleanup Kit (New England BioLabs, cat. number T1030), the 7.3 kb PCR product was transformed into *V. natriegens* strain TND1964 using the method described by Dalia *et al.*[1] Transformants were selected on LBv2 agar plates supplemented with ampicillin (25 μg/mL) and kanamycin (200 μg/mL). After screening by colony PCR, 18 out of 20 colonies were positive with T7pol-sc-F and T7pol-sc-R primers (Supplementary Figure I.2). Genomic DNA of these transformants were extracted and used as template in PCR with Vna-deldns-F1 and Vna-deldns-R2 primers. The resulting PCR products were sequenced with T7pol-seq-F1, T7pol-seq-F2, T7pol-seq-R1, T7pol-seq-R2, T7pol-seq-R3, araC-seq-F and araC-seq-R to verify the accuracy of the construct, allowing the identification of the strain V54 with correct sequence (sequencing results can be found in the Sequence Read Archive under accession numbers SRR32563228 and SRR32563229).

In the second round of MuGENT (multiplex genome editing by natural transformation), the strain V54 was subjected to the following genome edits: inactivation of *nadR* and *pncC,* two genes involved in the catabolism of NMN and introduction of FtNadE catalyzing the synthesis of NMN from nicotinic acid mononucleotides (NaMN).[2] To this end, the *pncC* gene was deleted by introducing the construct $P_{T7}$-FtNadE-$Sm^R$ into the strain V54. This construct was generated by Gibson assembly from the following fragments:

- the flanking left sequence (2.7 kb) which was amplified by PCR from *V. natriegens* genomic DNA in a 50 μL reaction containing 0.5 μM of Vna-delpncC-F1 and 0.5 μM Vna-delpncC-R1, 100 ng of extracted DNA, 0.2 mM dNTPs, and 0.02 U/μL of the Q5 Hot Start High-Fidelity DNA Polymerase in 1X Q5 Reaction Buffer. The thermocycling conditions of the PCR consisted of 98°C for 30 s, followed by 30 cycles of 98°C for 10 s, 65°C for 30 s, and 72°C for 1 min 30s.

- FtNadE coding sequence under the control of T7 promoter (0.8 kb) which was synthesized by Genscript. The fragment was amplified in a 50 μL reaction containing 0.5 μM of T7pro-FtNadE-gb-F and 0.5 μM FtNadE-gb-R, 1 ng of template, 0.2 mM dNTPs, and 0.02 U/μL of the Q5 Hot Start High-Fidelity DNA Polymerase in 1X Q5 Reaction Buffer. The thermocycling conditions of the PCR consisted of 98°C for 30 s, followed by 6 cycles of 98°C for 10 s, 58°C for 30 s, and 72°C for 30s and 9 cycles of 98°C for 10 s, 71°C for 1 min.

- $Sm^R$ fragment (0.9 kb) which was produced by PCR amplification in a 50 μL reaction containing 0.5 μM of SmR-gb-F and 0.5 SmR-gb-R, 1 ng of pCDFDuet™-1 vector, 0.2 mM dNTPs, and 0.02 U/μL of the Q5 Hot Start High-Fidelity DNA Polymerase in 1X Q5 Reaction Buffer. The thermocycling conditions of the PCR consisted of 98°C for 30 s, followed by 15 cycles of 98°C for 10 s, 63°C for 30 s, 72°C for 30 s.

- The flanking right sequence (2.4 kb) which was amplified by PCR from *V. natriegens* genomic DNA in a 50 μL reaction containing 0.5 μM of Vna-delpncC-F2 and 0.5 μM Vna-delpncC-R2, 100 ng of extracted DNA, 0.2 mM dNTPs, and 0.02 U/μL of the Q5 Hot Start High-

Fidelity DNA Polymerase in 1X Q5 Reaction Buffer. The thermocycling conditions of the PCR consisted of 98°C for 30 s, followed by 30 cycles of 98°C for 10 s, 67°C for 30 s, and 72°C for 1 min 20s.

Subsequently, these four PCR products were verified by agarose gel electrophoresis (Supplementary Figure I.3) and purified by Monarch® PCR & DNA Cleanup Kit (New England BioLabs, cat. number T1030). The fragments were then assembled by NEBuilder® HiFi DNA Assembly Master Mix (New England Biolabs, cat. number E2621L) and the resulting product of assembled reaction was amplified in a 50 µL reaction containing 0.5 µM of Vna-delpncC-F1 and 0.5 µM Vna-delpncC-R2, 1 µL of assembled product, 0.2 mM dNTPs, and 0.02 U/µL of Q5 Hot Start High-Fidelity DNA Polymerase in 1X Q5 Reaction Buffer. The thermocycling conditions consisted of an initial denaturation at 98°C for 30 s, followed by 15 cycles of 98°C for 10 s, 67°C for 30 s, and 72°C for 4 min. After verification by agarose gel electrophoresis (Supplementary Figure I.3), the 6.8 kb PCR product was purified by Monarch® PCR & DNA Cleanup Kit (New England BioLabs, cat. number T1030).

Besides, *nadR* gene was knock out by cotransformation with the construct $P_{T7}$-FtNadE-$Sm^R$ using a PCR product generated from the following fragments:

- the flanking left sequence (3.2 kb) which was amplified by PCR from *V. natriegens* genomic DNA in a 50 µL reaction containing 0.5 µM of Vna-delnadR-F1 and 0.5 µM Vna-delnadR-R1, 100 ng of extracted DNA, 0.2 mM dNTPs, and 0.02 U/µL of the Q5 Hot Start High-Fidelity DNA Polymerase in 1X Q5 Reaction Buffer. The thermocycling conditions of the PCR consisted of 98°C for 30 s, followed by 30 cycles of 98°C for 10 s, 69°C for 30 s, and 72°C for 2 min.

- the flanking right sequence (3.2 kb) which was amplified by PCR from *V. natriegens* genomic DNA in a 50 µL reaction containing 0.5 µM of Vna-delnadR-F1 and 0.5 µM Vna-delnadR-R1, 100 ng of extracted DNA, 0.2 mM dNTPs, and 0.02 U/µL of the Q5 Hot Start High-Fidelity DNA Polymerase in 1X Q5 Reaction Buffer. The thermocycling conditions of the PCR consisted of 98°C for 30 s, followed by 30 cycles of 98°C for 10 s, 70°C for 30 s, and 72°C for 2 min.

These two PCR products were verified by agarose gel electrophoresis (Supplementary Figure I.4) and purified by Monarch® PCR & DNA Cleanup Kit (New England BioLabs, cat. number T1030). The fragments were then assembled by NEBuilder® HiFi DNA Assembly Master Mix (New England Biolabs, cat. number E2621L) and the resulting product of assembled reaction was amplified in a 50 µL reaction containing 0.5 µM of Vna-delnadR-F1 and 0.5 µM Vna-delnadR-R2, 1 µL of assembled product, 0.2 mM dNTPs, and 0.02 U/µL of Q5 Hot Start High-Fidelity DNA Polymerase in 1X Q5 Reaction Buffer. The thermocycling conditions consisted of an initial denaturation at 98°C for 30 s, followed by 15 cycles of 98°C for 10 s, 69°C for 30 s, and 72°C for 4 min. After verification by agarose gel electrophoresis (Supplementary Figure I.4), the 6.4 kb PCR product was purified by Monarch® PCR & DNA Cleanup Kit (New England BioLabs, cat. number T1030) and mixed with the construct $P_{T7}$-FtNadE-$Sm^R$ for cotransformation into the strain V54 using the method described by Dalia *et al*.[1] Transformants were selected on LBv2 agar plates supplemented with ampicillin (25 µg/mL), kanamycin (200 µg/mL) and spectinomycin (500 µg/mL). After screening by colony PCR, only one out of 64 colonies was positive with nadR-sc-F and nadR-sc-R primers (Supplementary Figure I.5A). This strain, called V54-33, was also positive with T7pro-FtnadE-gb-F and FtnadE-gb-R primers (Supplementary Figure I.5B). Genomic DNA of V54-33 was extracted and used as template in PCR with Vna-delpncC-F1 and Vna-delpncC-R2. The resulting PCR product was sequenced with FtNadE-seq-F to confirm the accuracy of the construct (sequencing results can be found in the Sequence Read Archive under accession number SRR32563230).
