## Supplement II for "β-nicotinamide mononucleotide production in *Vibrio natriegens*: a preliminary study"

### SUPPLEMENT II
#### Generation of pACYCDuet-Nampt plasmids

**Supplementary Table II.1. Primers used for generation of pACYCDuet-Nampt plasmids.**

| Primer name | Primer sequence (5'→3') | Description |
|---|---|---|
| NamptCP-F | taagaaggagatatacatatg ACCAAGGAAAACCT | *Nampt* from *C. pinensis* |
| NamptCP-R | agcagcctaggttaatta GATAGTTGCGTTCTTG | |
| NamptSSC-F | taagaaggagatatacatatg AAGAACCTGATTCTTG | *Nampt* from *Sphingopyxis sp.* C-1 |
| NamptSSC-R | agcagcctaggttaatta GCGACCTTCTGAA | |
| NamptHD-F | taagaaggagatatacatatg GACAATCTTCTCAACT | *Nampt* from *H. ducreyi* |
| NamptHD-R | agcagcctaggttaatta CAGTGTGGTACGTG | |
| NamptVP-F | taagaaggagatatacatatg CTAAATCTTAATCAAAATATCG | *Nampt* from *Vibrio* phage KVP40 |
| NamptVP-R | agcagcctaggttaatta CGCAGTTTGAATCTTT | |
| pACYC-T7pro1-F | taattaacctaggctgct AGTCGAACAGAAAGTAATCGTATTGTACA | pACYCDuet™-1 linearization |
| pACYC-T7pro1-R | catatgtatatctccttctta TCCCCTATAGTGAGTCGTATTAATTTCCT | |
| nampt-sq-F | GGATCTCGACGCTCTCCCT | Sequencing primers |
| nampt-sq-R | GATTATGCGGCCGTGTACAA | |

**Lower case nucleotides specify overlap regions*

***>NamptCP (Nampt from Chitinophaga pinensis)***

ATGACCAAGGAAAACCTGATCTTGTTAGCAGACGCGTACAAGTATAGCCATCACAAGCTGTACATTCCTGGTACCGAGTATATCTACAGCTACTTCGA
GTCACGTGGTGGCAAATTCAACGAAACGGTGTTTTACGGTTTGCAGTATTTCCTGATGGAATACTTGCAGGGTGCAGTCATTACCAAAGAGAAGCTGG
ATGAAGCGGAGGCAACGTTACTGGAAGTGTTCGGTCGCAACGACGTCTTTGATCGTACTCGCTTCGAGTACATTATCGAAAAGCACGGTGGACGTTTA
CCGGTTCGCATCAAAGCAGTGCCAGAAGGTACTGTCACCGGAGTTCGTAACGTGCTGATGACGATTGAGAATACTGATCCTAACTGTTACTGGGTTAC
CAACTTCCTGGAAACGTTGCTTATGCAGATTTGGTATCCGTGTACTGTGGCGACTCTGTCACGTGAAATCAAGAAAACCGTCAAGCAGTACTATAACG
AGACAGCAAGCGAAGCTGCGTTTGCAGGTATTGATTTCGTTTTGAACGACTTTGGCTTCCGTGGTGCTAGTAGCGTTGAATCAGCAGGAATTGGTGGC
TCTGCTCATCTGATTAACTTCAGTGGCTCAGATACCTTGATCGGTAGCACTTTTGCAAAACGCTATTACCAAGCTCCTGTTGCACCAGGCTTAAGCAT
TCCGGCAACTGAACACTCAATCGTGACAATGCTGGGTGAGGAAGGCGAGTTAGAAATCTTTCGCCATATTCTGAATGCGTTCCCTACCGGTACTATCG
CATGTGTTAGCGATAGTTACAACATTCTGCGTGCTTGTCGCGAGTATTGGGGAACTGAATTGAAAGAGCAGATCTTATCACGCCAAGGTACACTGGTT
ATTCGTCCGGATAGTGGCGACGCGATTCAGACCTTGCTGAAAGTGTTTGAGATCCTGATGGAAACGTTCGGTTATACTGTCAATGAGAAAGGCTACAA
GGTTTTACCTCCACAAGTGCGTGTCATTCAGGGTGATGGAATCAGCTATTCTTCAATTCCTCCGATCTTCGAAGCACTGAAGCAAGCTGGTATTAGTG
CAGAGAATCTTGTGTTAGGCATGGGAGGTGCTTTGCTGCAACGTGTTAACCGCGATACCCAGGAATACGCACTGAAATGTAGCTTTGCGCAAGTGAAT
GGTAAGGCTATTAACGTACAGAAAAACCCTTTGGAGTTAGATGCGAATGGTAACACTCGCGTCTCATTCAAAAAGTCGAAAAGCGGTAAGCAGAAACT
GGTGGTTGAAAACGGCATCTATACTAGTTTACCGGAGAATGAAGCACCAGCTCTTGCGGATCAACTGGTTACCGTGTTCGAAGACGGAGAGATCAAGA
AAGCGTACAGCTTTGAACAGATTCGCAAGAACGCAACTATCTAA

***>NamptHD*** **(*Nampt from Haemophilus ducreyi*)**

ATGGACAATCTTCTCAACTATAGCAGTCGTGCTAGTGCCATTCCGTCTTTGTTATGTGATTTCTACAAGACATCACATCGCATCATGTATCCAGAGTG
TTCGCAGATTATCTACTCCACGTTTACTCCTCGTAGCAATGAACAAGCACCGTATCTAACCCAGGTCGTGTCATTCGGATTTCAGGCCTTCATTATCA
AATACCTGATACACTATTTCAATGACAACTTTTTCAGTCGCGATAAACACGATGTTGTAACTGAGTATTCGGCATTCATCGAGAAGACATTGCAACTC
GAAGACACCGGTGAGCATATTGCTAAACTCCACGAACTTGGCTACCTACCAATTCGTATCAAGGCGATTCCTGAAGGCAAAACGGTTGCCATCAAGGT
GCCGGTAATGACAATTGAGAACACTCATAGCGACTTTTTCTGGTTGACGAACTATCTGGAAACCCTCATCAATGTCTCTTTATGGCAACCAATGACTA
GTGCCTCAATTGCGTTTGCATACCGCACAGCTCTAATCAAATTCGCAAATGAGACGTGTGACAACCAGGAACACGTTCCTTTTCAGAGCCACGATTTC
AGTATGCGTGGAATGTCTTCACTGGAGTCGGCTGAAACCAGTGGTGCAGGGCATTTGACGAGCTTCCTCGGCACCGATACAATTCCAGCCTTATCGTT
TGTAGAGGCTTACTATGGATCTAGTTCACTTATCGGTACTTCGATTCCTGCGTCAGAACACTCTGTGATGAGCTCTCATGGCGTCGACGAACTAAGTA
CGTTTCGCTATCTGATGGCTAAGTTTCCGCACAATATGTTGTCGATCGTATCTGACACTACCGACTTCTGGCACAATATCACAGTGAACTTACCACTC
CTTAAGCAAGAGATCATTGCACGCCCTGAAAATGCCCGATTAGTCATCCGTCCAGATTCAGGAAACTTTTTCGCGATCATTTGTGGTGATCCGACTGC
CGATACGGAGCATGAACGCAAAGGCCTAATCGAGTGTCTTTGGGACATCTTTGGAGGTACCGTTAATCAGAAAGGCTATAAGGTGATCAATCCTCACA
TTGGAGCTATCTATGGCGATGGTGTAACATACGAAAAGATGTTCAAAATCCTCGAGGGCTTACAAGCTAAAGGATTTGCATCTAGCAACATCGTGTTC
GGTGTTGGAGCCCAAACGTATCAGCGCAATACTCGTGATACCTTAGGCTTTGCGTTGAAGGCTACAAGTATAACTATTAATGGAGAAGAGAAAGCAAT
CTTCAAGAACCCAAAAACCGATGACGGATTTAAGAAAAGCCAGAAGGGTCGTGTCAAGGTACTGAGTCGCGACACGTACGTAGATGGCTTGACATCTG
CAGACGATTTCTCGGATGACTTACTCGAGCTGTTGTTTGAGGATGGTAAACTACTTCGTCAAACTGACTTCGACGAAATCCGCCAGAATCTCCTTGTC
TCACGTACCACACTGTAA

***>NamptSSC (Nampt from Sphingopyxis sp. C-1)***

ATGAAGAACCTGATTCTTGCGACCGATAGCTACAAACAGTCACACTTCTTGCAATATCCGCCTGAAGCACGCGTTATCAGTGCGTATGTGGAGGCACG
TCCAAATCCGTTTTCTGAAGAGATTGTGTTCCTGGGTCTTCAGCCTTTACTGGTTGATTACTTTAGCCAACCGATTAACGCAGCGGACATCGATGAAG
CAGAGGCTATTTGTATCGCCCATGGAGTTCCATTCAATCGTGCAGGTTGGGAAGCGATTGTGGCAGATCATGGAGGCTATCTGCCTCTGGAGATCAAA
GCCTTGCCGGAAGGTGCAATTGTTCCTGCCGGTGTACCGTTAGTGCAGTTGGAAAACACTGATCCACGCATGCCTTGGCTTACCACGTTTATCGAGAC
TGCGATGTTACGTGCAATTTGGTATCCAACCACAGTTGCCACTCTGTCGTGGAAGTGTAAACAGGTAATCCGTGCTGGCTTAGAGAAGACCTCAGACG
ATGTCGAAGGTCAGTTGCCGTTCAAGCTTCACGACTTTGGTGCACGTGGAGTTAGTAGCGCTGAATCAGCAGGTTTAGGCGGACTGGCACATCTTGTG
AACTTCCAAGGCACTGATACGATGGAGGCGTTGGTTGCTGCACGTCGCTACTATGGTGCCGATATGGCTGGCTTCAGTATTCCTGCAGCGGAACATTC
AACCATGACTAGCTGGGGACGCGATCGTGAAGAGGACGCGTATCGCAATATGTTGGATCGTTTTGAAGGCGAGGGTCGCATTGTTGCCGTGGTCAGCG
ACAGTTACGACCTTGATACTGCGGTAACCGATATCTGGGGTGGCTCACTGCGTGAAAAGGTGTTAGGTCGTGCAGGCACCTTGGTGGTTCGTCCAGAT
AGTGGTGACCCTATTGAGACTCCGTTACGCACGGTTAAGACACTGTGGGAAAAGTTCGGTGGACATGTCAACGGCAAAGGTTATCGCGTGTTGGATCC
TCATGTTCGTGTAATCCAAGGCGATGGAATGACCGTAGACTCAATTGGTCGCTTAGTGCAGCGTATGATCGAAGAGGGCTTTGCAATCGACAACATTG
CGTTCGGCATGGGAGGTGGCATGCTTCAGCATGTTAACCGTGATACCTTACGCTTTGCCATGAAAGCGAATGCTATGTTGGGCAGTGATGGTGTCTGG
CACGACGTGTTCAAGATGCCGTCAACCGATCCTGGCAAAGCTAGCAAGGCAGGTCGTCAAGCCGTTGTACTGAAAGACGGACGTATGGCTGCGGCACG
TTTAGATAGTGTGGCAGTAGGTGAAGATCTTCTGGTTCCTGTCTGGCGTAATGGCGAATTACTGGTTCGCCATGATTTCGACGCTGTGCGCAAACGTT
CAGAAGGTCGCTAA

***>NamptVP (Nampt from Vibrio phage*** **KVP40*)***

ATGCTAAATCTTAATCAAAATATCGCAATCGCAACTGACAGCTACAAAGTTTCACACTGGTCTCAATTCCCTCGTGGTCTAGAGTACAGCCAATATTA
CGTCGAAAGCCGTGGTGGTAAGTTCGACAAAATTATGGTCGATGGTATGGCATACATGTGTCGTATTCTCGAAAAAGGTGTTTCGATGAACGATGTCA
AGCGTGCGAAACGTCTGTTCAAAAAGCACTTCGGTTCTGAAGTCTTCAATGACAAGGGTTGGGACATCATCGTTAACGAACTGAAAGGCAAACTGCCT
ATTAAGATTCGTGCGGTCAAAGAAGGTACTGTAGTACCAGTTAAACACCCTATCTTGACGATTGAGAACACAGATCCGCGTTTTGGTTGGCTGCCTGG
CTATCTTGAAACGTTCATCTTGCGTGCTCTTTGGTACCCGACAACTGTAGCAACGATCTCGTTCGAAGTGAAGAAAATCATTCGCCAATTCATGAAGA
AGACGGTTGATGATGAACGCATTGCAGAACAAGAGCCGTTCAAACTTCACGACTTTGGTTCTCGTGGTGTATCAAGTGGCGAATCTGCGGCTATCGGC
GGCTCTGCTCACTTGAAAAACTTCTTGGGTACTGACACTGTAGAAGCACTTGTTGCAGTTGAAGAACTTTACGCTGAAGATGTTGAAGACTTCATTGC
AGGATTCTCAATCCCTGCACGTGAACACTCAACAACTACAATCTACAAAGAAGCTGGTGAAGACCAAGCATTCTTGAACTCTATTGAGCAATGGGGTG
CTGCATTATACGCTTGTGTAATGGACAGCTACGACTACGAAGCAGCAATGAATCGTGTCTCGACTGGTCGCTTCAAAGAGCTAATCATCAGCAAAGGC
GGTACGTTTGTTGCTCGTCCTGACTCTGGTGTGCCTGTAGATGTAGTAATGAAAGGTCTTGAGATTCTTGGTAAAAACGTTGGTTATACAATCAACAG
CAAGGGTTACAAAGTATTGCACCCTAGCTACCGTATCATTCAAGGTGACGGCGTGAACATCGAAGAAATTCGTCGAATCTTGTCTTACATGGAAAGTA
AAGGCTGGTCGGCAGAAAATATTGCATTCGGTATGGGTGGCGGGTTGCTTCAACAACTTGACCGTGATACTCAACGCTTTGCAATGAAAATGTCAGCG
GCAATCATCAACGGCGAATACGTTAGCGTGTTCAAAATGCCGAAGACTGATCCGACAAAAGCATCAAAAGCAGGTTTCTTGGATTTGATTGCAGTTGA
TGCAGACAATCCGAACCCAGCGGCTCGTGGTTACGTGACGTTCTCTTCTGAAGATTACGACAATCGCGTACACCCGAAAAGCGTGATGCAGACAATCT
TCGAAGACGGTGTAACAGTTGCCGACTTCTCACTAGAAGAAGCGCGTAAACTGTCTGACGTACAAGCTGATTTCCTGAACGAAGGTGAATGGAAAATC
GCGAAAAAGATTCAAACTGCGTAA

**Supplementary Figure II.1. Codon-optimized sequences encoding NAMPT from *Chitinophaga pinensis*, *Haemophilus ducreyi*, *Sphingopyxis* sp. C-1 and *Vibrio* phage KVP40**

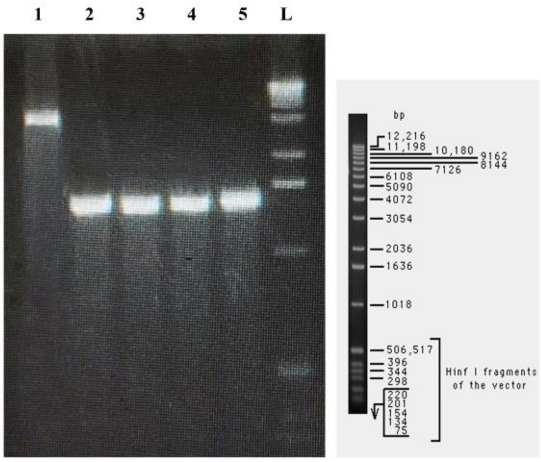


**Supplementary Figure II.2. PCR products of the fragments used for generating *pACYCDuet-Nampt plasmids* by Gibson assembly.**

*(1) pACYCDuet-1 vector linearized with pACYC-T7pro1-F and pACYC-T7pro1-R primers; (2) Nampt from Chitinophaga pinensis; (3) Nampt from Sphingopyxis sp. C-1; (4) Nampt from Haemophilus ducreyi; (5) Nampt from Vibrio phage KVP40; L: 1 kb DNA ladder (Invitrogen)*

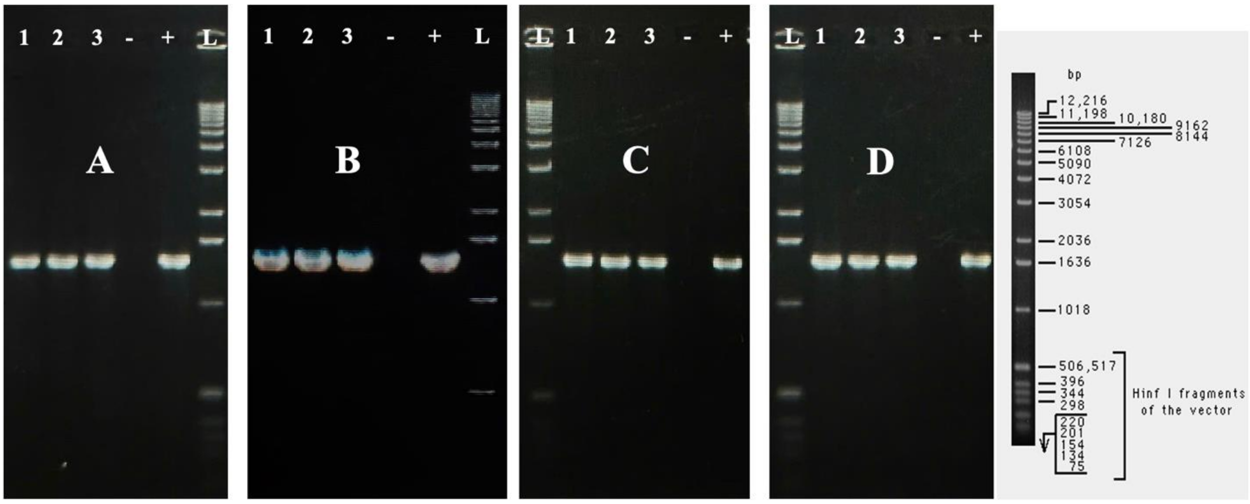


**Supplementary Figure II.3. Screening results for transformants harboring pACYCDuet-Nampt from *Chitinophaga pinensis* (A), *Sphingopyxis* sp. C-1 (B), *Haemophilus ducreyi* (C), and *Vibrio* phage KVP40 (D).**

*(-) no-template control; (+) positive control; 1-3: colony PCR products; L: 1 kb DNA ladder (Invitrogen).*

In this study, codon-optimized sequences encoding NAMPT (Supplementary Figure II.1) were synthesized by Genscript. These sequences were amplified by PCR in 50 µL-reactions containing 0.5 µM of specific pairs of primers (each) (Supplementary Table II.1), 1 ng of template, 0.2 mM dNTPs and 0.02 U/µL of the Q5 Hot Start High-Fidelity DNA Polymerase in 1X Q5 Reaction Buffer. The thermocycling conditions of the PCR consisted of 98°C for 30 s, followed by 15 cycles of 98°C for 10 s, 57°C for 30 s and 72°C for 45 s.

The vector pACYCDuet™-1 was linearized by PCR in a 50 µL reaction containing 0.5 µM of pACYC-T7pro1-F and pACYC-T7pro1-R primers (each), 0.1 ng of vector, 0.2 mM dNTPs and 0.02 U/µL of the Q5 Hot Start High-Fidelity DNA Polymerase in 1X Q5 Reaction Buffer. The thermocycling conditions consisted of 98°C for 30 s, followed by 15 cycles of 98°C for 10 s, 66°C for 30 s, 72°C for 2 min. The resulting 4047 bp PCR product was *Dpn*I treated to remove the template.

Subsequently, all PCR products were verified by agarose gel electrophoresis (Supplementary Figure II.2) and purified by Monarch® PCR & DNA Cleanup Kit (New England BioLabs, cat. number T1030). These two fragments were assembled by NEBuilder® HiFi DNA Assembly Master Mix (New England Biolabs, cat. number E2621L) according to the manufacturer's instructions. The resulting products were transformed into *E. coli* DH5α Competent Cells by the heat shock method. Transformants were selected on LB agar plates supplemented with Chloramphenicol (35 µg/mL). After colony PCR screening with specific pairs of primers for each *Nampt* gene (Supplementary Figure II.3), plasmids were extracted by Monarch® Plasmid Miniprep Kit (New England BioLabs, cat. number T1010) and sequenced with nampt-sq-F and nampt-sq-R to verify the accuracy of the constructs. Sequencing results of the constructs can be found in the Sequence Read Archive under accession SRR32563234, SRR32563232, SRR32563233 and SRR32563231.
