## Supplement III for "β-nicotinamide mononucleotide production in *Vibrio natriegens*: a preliminary study"

### SUPPLEMENT III
#### Quantification of intracellular NMN concentration by HPLC

**Supplementary Table III.1. Quantification of intracellular NMN concentration by HPLC**

| Sample | NMN Concentration by HPLC (mg/L extract) | Dilution factor | Intracellular NMN Concentration (mg/L culture) | Intracellular NMN Concentration (µM) |
|---|---|---|---|---|
| CP1* | 0.88 | 18.1 | 15.93 | 47.7 |
| CP2* | 0.90 | 15.3 | 13.77 | 41.2 |
| HD1* | 0.28 | 15.8 | 4.42 | 13.2 |
| HD2* | 0.27 | 13.5 | 3.65 | 10.9 |
| SSC1* | < LOD** | NA | NA | NA |
| SSC2* | < LOD** | NA | NA | NA |
| VP1* | 0.55 | 16.5 | 9.08 | 27.2 |
| VP2* | 0.34 | 18.9 | 6.43 | 19.3 |
| V54-33.1 | < LOD** | NA | NA | NA |
| V54-33.2 | < LOD** | NA | NA | NA |

*CP: *Chitinophaga pinensis,* HD: *Haemophilus ducreyi,* SSC: *Sphingopyxis sp.* C-1, VP: *Vibrio Phage* KVP40

**LOD = 0.04 mg/L

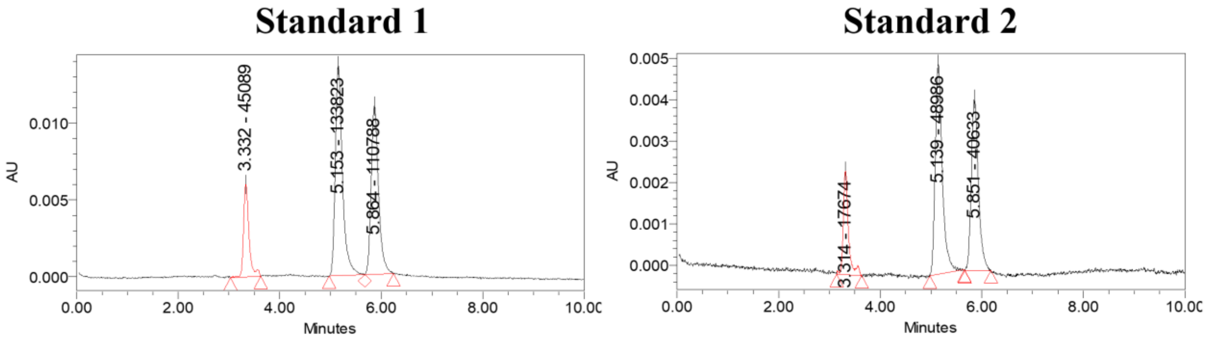


**Supplementary Figure III.1. HPLC chromatograms of NMN standards (retention times around 3.3 minutes)**

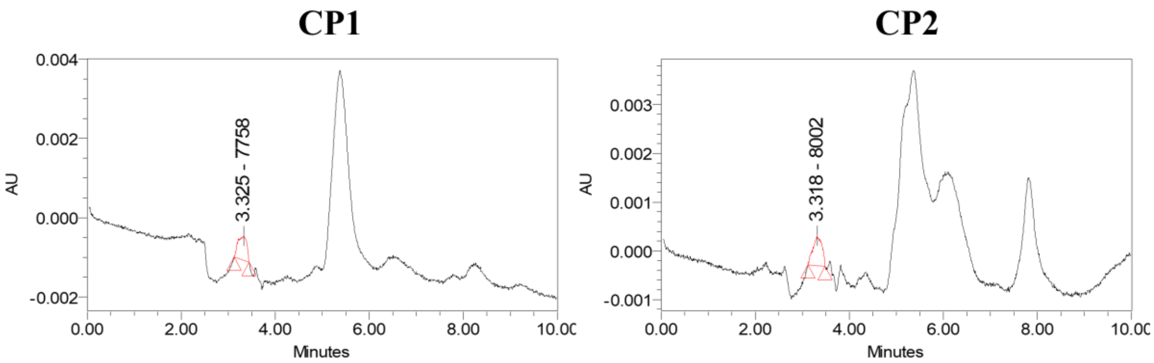


**Supplementary Figure III.2. HPLC chromatograms of transformants harboring pACYC-Nampt-CP**

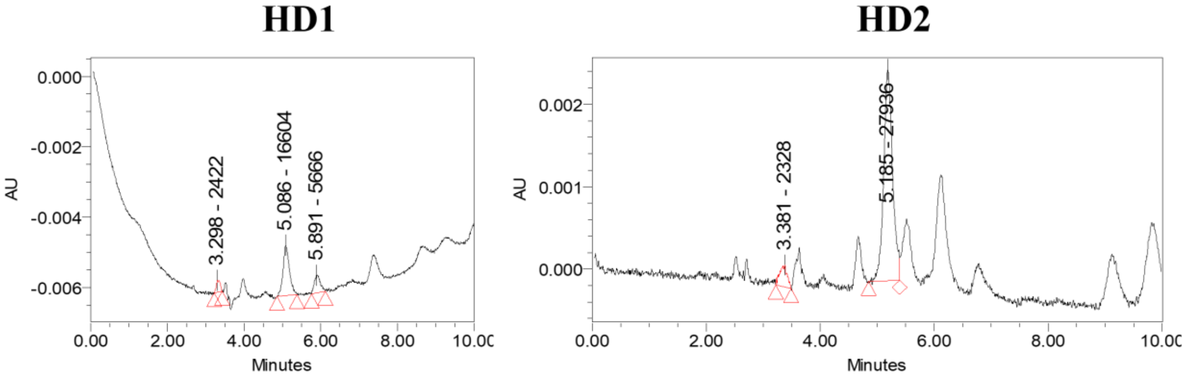


**Supplementary Figure III.3. HPLC chromatograms of transformants harboring pACYC-Nampt-HD**

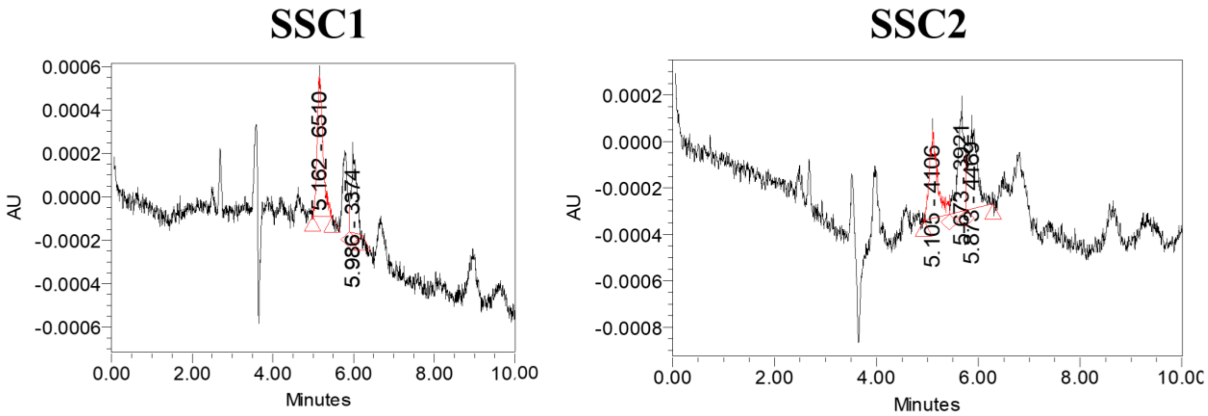


**Supplementary Figure III.4. HPLC chromatograms of transformants harboring pACYC-Nampt-SSC**

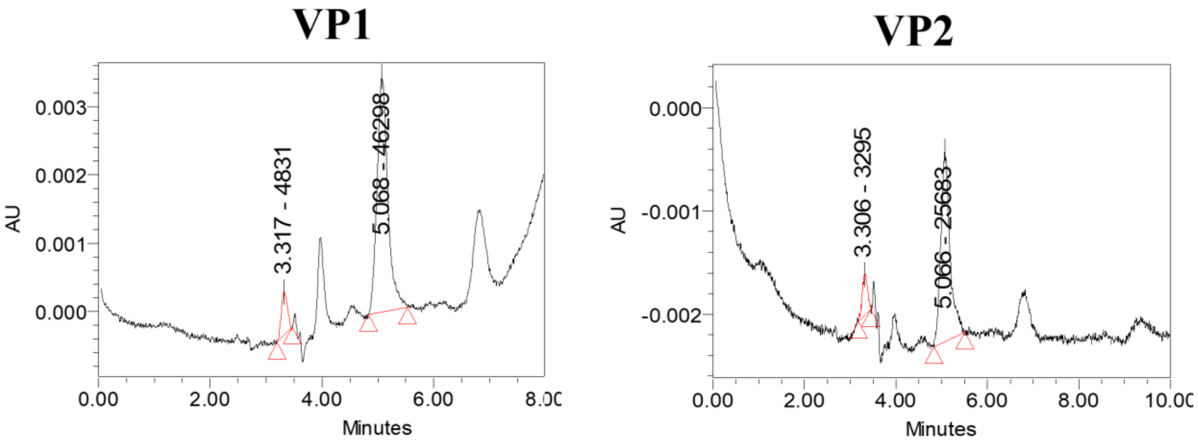


**Supplementary Figure III.5. HPLC chromatograms of transformants harboring pACYC-Nampt-VP**

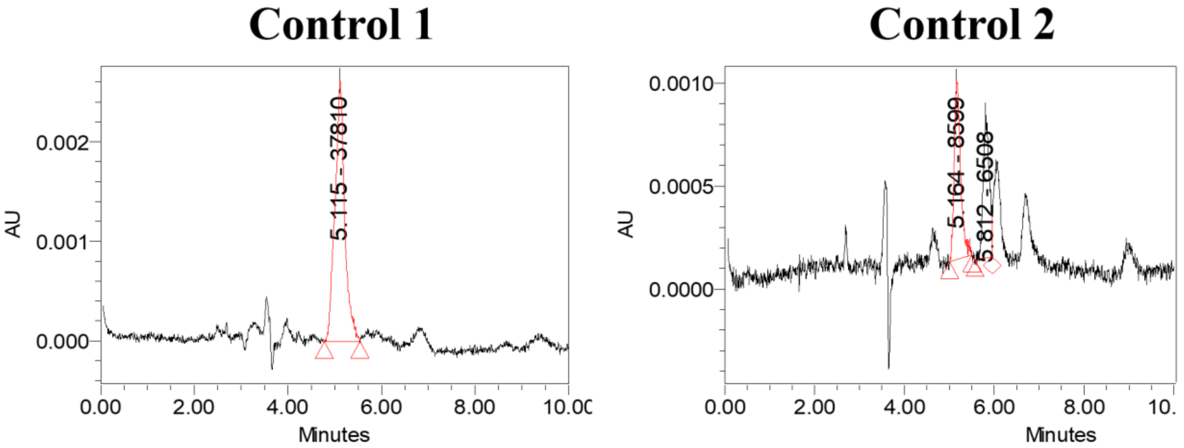


**Supplementary Figure III.6. HPLC chromatograms of V54-33 strain**
